# The segmented flavivirus Alongshan virus reduces mitochondrial mass via degrading STAT2 to suppress innate immune response

**DOI:** 10.1101/2024.03.06.583679

**Authors:** Yinghua Zhao, Liyan Sui, Mingming Pan, Fangyu Jin, Yuan Huang, Shu Fang, Mengmeng Wang, Lihe Che, Wenbo Xu, Ning Liu, Nan Liu, Haicheng Gao, Zhijun Hou, Fang Du, Zhengkai Wei, Jixue Zhao, Kaiyu Zhang, Yicheng Zhao, Ze-Dong Wang, Quan Liu

**Author notes:** Correspondence to Quan Liu.

## Abstract

Alongshan virus (ALSV) is a newly discovered pathogen of the *Flaviviridae* family, characterized by a multi-segmented genome distantly related to the canonical flaviviruses. Understanding the pathogenic mechanism of the emerging segmented flavivirus is crucial for the development of intervention strategies. Here we show that ALSV can infect multiple mammalian cells and induces the expression of antiviral genes. Moreover, ALSV is sensitive to IFN-β and possesses the ability to counteract type I IFN response. Mechanistically, ALSV’s nonstructural protein NSP1 binds to and degrades human STAT2 through an autophagy pathway in a species-dependent manner, resulting in direct inhibition of the expression of interferon-stimulated genes (ISGs). Specifically, NSP1 methyltransferase domain binds to the key sites of F175/R176 located in coiled-coil domain of STAT2. Moreover, NSP1-mediated degradation of STAT2 results in a reduction in mitochondrial mass by disrupting mitochondrial dynamics to induce mitophagy and inhibiting mitochondrial biogenesis, thereby suppressing the innate immune response. Interestingly, inhibiting mitophagy using 3-Methyladenine and enhancing mitochondrial biogenesis using the PPARγ agonist pioglitazone can reverse NSP1-mediated inhibition of ISGs, suggesting that promoting mitochondrial mass presents an effective antiviral strategy. Our findings elucidate the intricate regulatory crosstalk between ALSV and the host’s innate immune response, providing valuable insights into the pathogenesis and intervention strategy of emerging segmented flavivirus.

## Introduction

In recent decades, a diverse range of vector-borne viruses capable of infecting humans and causing diseases has come to light (Madison-Antenucci et al., 2020; Ni et al., 2023). Alongshan virus (ALSV), a segmented flavivirus in the *Flaviviridae* family, was identified in patients who had experienced tick bites in Northeastern China (Wang et al., 2019a; Wang et al., 2019b). Subsequent researches have revealed the widespread presence of ALSV in various regions, including Russia, Finland, Switzerland, and Germany (Ebert et al., 2023; Kholodilov et al., 2021; Kholodilov et al., 2020; Kuivanen et al., 2019; Stegmüller et al., 2023). ALSV infection leads to a condition known as Alongshan fever, characterized by common clinical symptoms such as fever, headache, skin rash, myalgia, arthralgia, depression, and coma (Wang et al., 2019a; Wang et al., 2019b; Zhang et al., 2020). Importantly, Jingmen tick virus (JMTV), another segmented flavivirus, also demonstrated the ability to infect humans, underscoring the emerging threat posed by segmented flaviviruses to human health (Jia et al., 2019; Qin et al., 2014; Yu et al., 2020b).

Several members in the *Flavivirus* genus are significant for human health, including dengue virus (DENV), West Nile virus (WNV), Zika virus (ZIKV), and yellow fever virus (YFV). These viruses share common characteristics as they are enveloped and carry a positive-sense single-stranded RNA genome, which consist of a single open reading frame (ORF) that encodes three structural proteins (capsid, membrane, and envelope) and seven nonstructural (NS) proteins (NS1, NS2A, NS2B, NS3, NS4A, NS4B, and NS5) (Barrows et al., 2018; Chambers et al., 1990; Mukhopadhyay et al., 2005). However, ALSV and JMTV represent distinct members of *Flaviviridae* family, as their genome comprises four single-stranded positive-sense RNA segments. Among these, segments 2 and 4 encode structural proteins VP1-VP4, the evolutionary origin of which remains unknown. Conversely, segments 1 and 3 encode the non-structural proteins NSP1 and NSP2, which individually exhibit homology to the NS5 and NS2B-NS3 of the mono-segmented flaviviruses (Gao et al., 2020; Wang et al., 2019a; Zhao et al., 2022). Notable, ALSV NSP1 has enzymatic activities of the methyltransferase (MTase) and the RNA-dependent RNA polymerase (RdRp), and play a critical role in viral replication (Chen et al., 2023). Nevertheless, the precise roles of NSP1 in regulating host response remain a topic of ongoing investigation, with its implications for both viral pathogenesis and host defense yet to be fully elucidated.

Innate immune response, particularly the type I interferons (IFN-I), play a crucial role in the host’s defense against viral infections (McNab et al., 2015). The signaling cascade mediated by IFN-I includes two key stages: IFN-I production and signal transduction. In the initial stage, the rapid production of IFN-I is triggered by the recognition of viral pathogen-associated molecular patterns (PAMPs) through host pattern recognition receptors (PRRs) (Iwasaki, 2012). In the second stage, the secreted IFN-α/β engages with the IFN-I receptors (IFNARs) to activate JAK/STATs cascades. The activated STATs combine with interferon regulatory factor 9 (IRF9) to create the transcription complex of IFN-stimulated gene factor 3 (ISGF3). ISGF3 then translocates into the nucleus, where it binds to interferon stimulated response elements (ISREs) and drives the expression of interferon-stimulated genes (ISGs) to suppress virus replication (Schneider et al., 2014). It is well-established that all known flaviviruses have evolved sophisticated strategies to antagonize IFN-I response, enabling them to infect vertebrate hosts successfully. For example, the NS5 proteins of DENV and ZIKV are known to degrade STAT2 to inhibit IFN-I response (Ashour et al., 2009; Ashour et al., 2010; Grant et al., 2016; Morrison et al., 2013). Here, we demonstrate that ALSV displays sensitivity to IFN-β and possesses the capability to counteract IFN-I response through NSP1. In a nutshell, ALSV’s NSP1 selectively targets human STAT2 for degradation, while having no effect on mouse STAT2. This degradation subsequently leads to a reduction in mitochondrial mass by disrupting mitochondrial dynamics and inhibiting mitochondrial biogenesis, resulting in the inhibition of innate immune response.

## Results

### ALSV infects multiple mammalian cells and induces antiviral response

As a newly discovered zoonotic pathogen, we investigated ALSV’s potential to infect mammalian cells. HEK293T cells, derived from human embryonic kidney, were infected with escalating doses of ALSV. At 48 hours post-infection (hpi), we examined viral replication levels which showed that the ALSV copies in supernatants and the ALSV segment 2 (*S2*) mRNA level in cells increased with the escalating amount of infectious virus (Fig. 1, A and B). In contrast, when HEK293T cells were infected with ALSV and incubated for 3, 6, 12, 24, and 48 hours (h), we observed that the ALSV copies in supernatants did not increase with the prolonged infection time (Fig. 1 C), and the *S2* mRNA level showed a downward trend with the extension of infection time (Fig. 1 D), suggesting that ALSV’s replication ability in HEK293T cells may be limit. Furthermore, we extended our investigation to human hepatoblastoma-derived HepG2 cells and African green monkey kidney-derived Vero cells. The temporal ALSV replication dynamics revealed that viral copies in HepG2-culture supernatants increased during the first 24 hpi and decreased at 48 hpi (Fig. 1 E), while the *S2* mRNA levels in HepG2 cells remained unchanged over time (Fig. 1 F), which is similar with that in Vero cells (Fig. 1, G and H). In summary, our findings indicate that ALSV can infect multiple mammalian cells, but exhibits limited replication capability.

**Figure 1.**
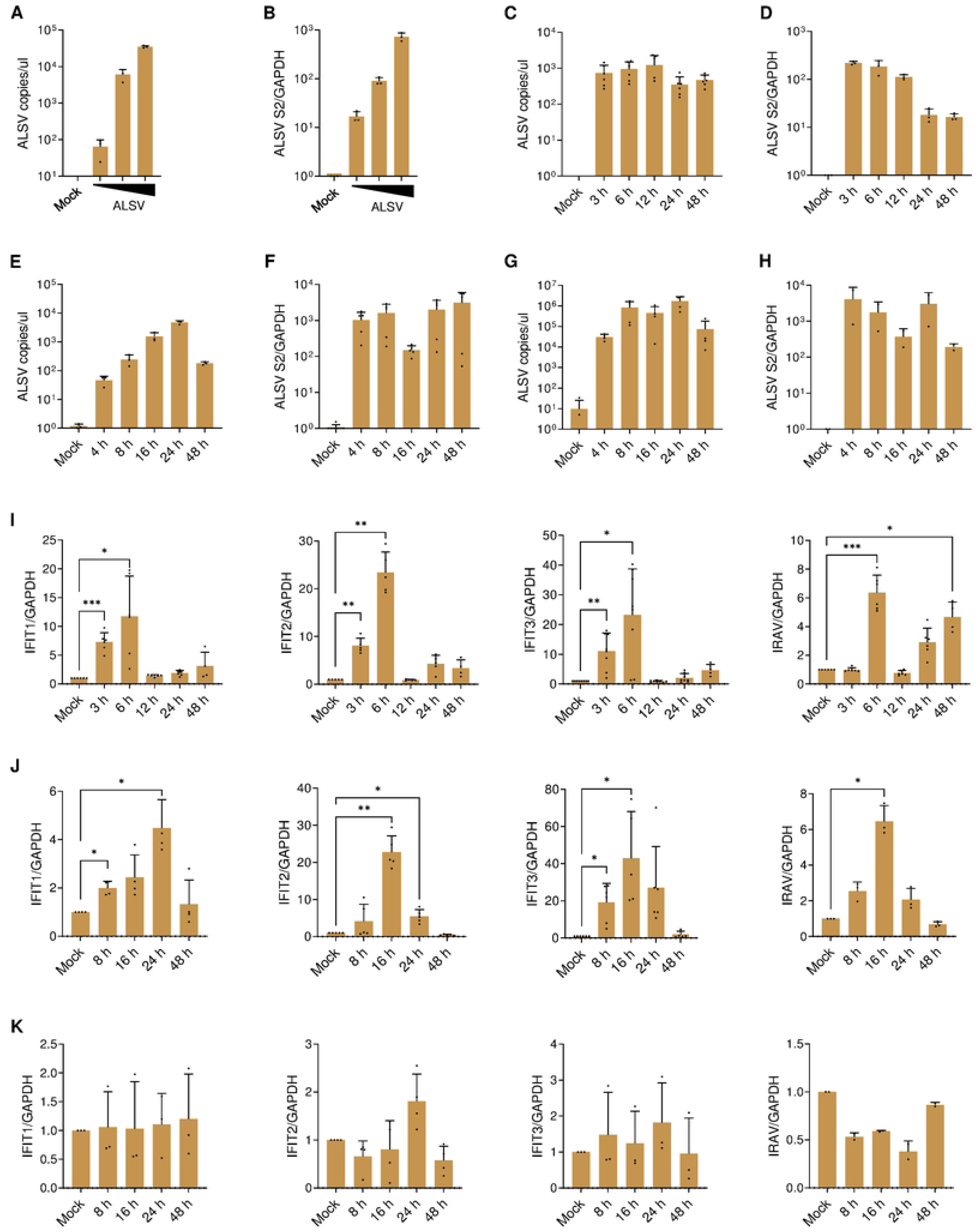
ALSV infection and host response in different cell types. **(A and B)** HEK293T cells were infected with escalating doses of ALSV (1 μl, 10 μl, and 100 μl). At 48 hours post-infection (hpi), viral RNA copies in supernatants were quantified using TaqMan-qPCR (A), and the mRNA levels of viral segment 2 (*S2*) relative to *GAPDH* control in cells were measured by qPCR (B). **(C and D)** HEK293T cells infected with ALSV were analyzed at various time points (3, 6, 12, 24, and 48 hpi). Viral RNA copies in supernatants were assessed by TaqMan-qPCR (C), and the mRNA levels of viral *S2* in cells were determined by qPCR (D). **(E and F)** HepG2 cells were infected with ALSV and analyzed at 4, 8, 16, 24, and 48 hpi for viral RNA copies in supernatants (E) and viral *S2* mRNA levels in cells (F). **(G and H)** Vero cells were infected with ALSV and assessed at 4, 8, 16, 24, and 48 hpi for viral RNA copies in supernatants (G) and viral *S2* mRNA levels in cells (H). **(I-K)** HEK293T (I), HepG2 (J), and Vero (K) cells were infected with ALSV, and the mRNA levels of host *IFIT1*, *IFIT2*, *IFIT3*, and *IRAV* genes in cells were measured by qPCR at the indicated time points, with GAPDH serving as the internal reference control. Statistical analysis was performed using one-way ANOVA with multiple comparison correction (\**P* < 0.05, \*\**P* < 0.01, and \*\*\**P* < 0.001).

ALSV exhibits more robust replication in IFN-I-deficient Vero cells compared to HEK293T and HepG2 cells (Fig. 1, C-H), suggesting that IFN-I may play a critical role in host’s antiviral response. To gain insights into the interaction between ALSV and the IFN-I system, we firstly examined whether ALSV infection induces the expression of ISGs, and found that ALSV infection stimulated the expression of *IFIT1*, *IFIT2*, and *IFIT3* as early as 3 hpi and reached its peak at 6 hpi, while *IRAV* peaked at 6 hpi and then recovered at 24 and 48 hpi in HEK293T cells (Fig. 1 I). In HepG2 cells, ALSV induced the expression of *IFIT1*, *IFIT2*, *IFIT3* and *IRAV* as early as 8 hpi, with peaks at 16 or 24 hpi, which occurred later than that in HEK293T cells (Fig. 1 J). This delay in ISGs expression might explain the increased ALSV copies in HepG2 cells (Fig. 1, C-F). Notably, ALSV did not stimulate ISGs expression in Vero cells (Fig. 1 K), which aligns with the fact that Vero cells are unable to produce IFN-I. These findings suggest that ALSV infection induces the expression of antiviral genes during the early stages of infection, contributing to the control of viral replication.

### ALSV exhibits sensitivity to IFN-β and antagonism to type I IFN signaling

To investigate ALSV’s sensitivity to IFN-I, we infected various cells with ALSV and treated them with the increasing concentrations of IFN-β for 24 h. The virus levels assay showed that IFN-β inhibited ALSV replication in a concentration-dependent manner (Fig. 2, A-C). Immunofluorescence assay (IFA) of viral dsRNA further confirmed that IFN-β effectively suppressed ALSV replication (Fig. 2 D). Moreover, we observed that the longer durations of IFN-β treatment led to more effective inhibiting of ALSV replication (Fig. 2 E). Overall, these data underscore the sensitivity of ALSV to the action of IFN-β.

**Figure 2.**
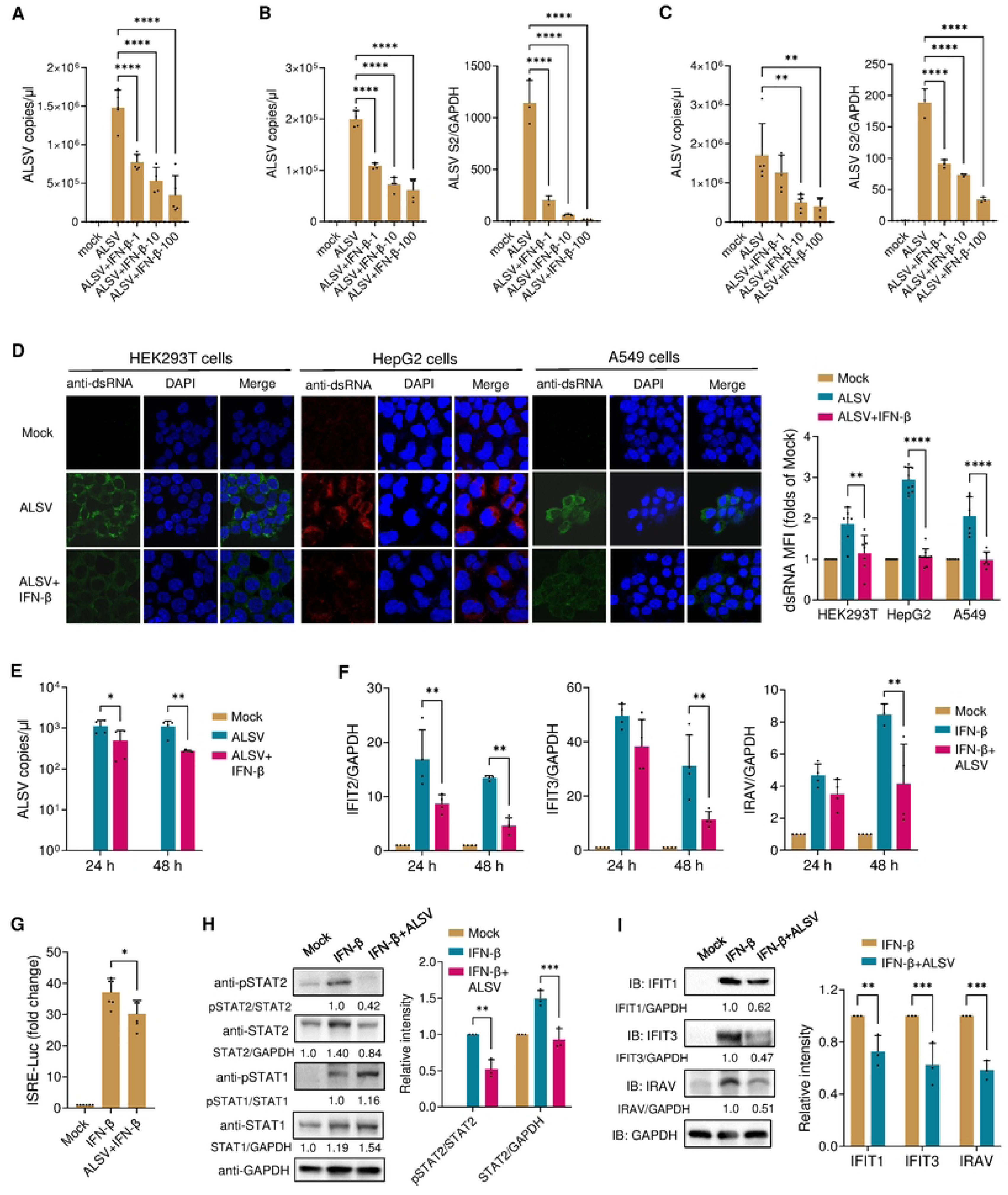
Sensitivity of ALSV to IFN-β and antagonism of type I IFN signaling. **(A-C)** HEK293T (A), HepG2 (B), and A549 (C) cells were infected with ALSV. At 2 hours post-infection (hpi), cells were treated with increasing concentrations of IFN-β (1, 10, 100 ng/ml) for 24 h. Viral copies in supernatant and the mRNA levels of viral *S2* in cells were determined using qPCR. **(D)** HEK293T, HepG2, and A549 cells were infected with ALSV. At 2 hpi, cells were treated with IFN-β (10 ng/ml) for 24 h and subjected to immunofluorescence analysis using an anti-dsRNA antibody. Nuclei were counterstained with DAPI. The mean fluorescence intensity (MFI) per graph of dsRNA was analyzed (n ≥ 6 graphs). **(E)** HepG2 cells were infected with ALSV. At 2 hpi, cells were treated with or without IFN-β for 24 or 48 h, and viral copies in supernatant were determined. **(F)** HepG2 cells treated with or without IFN-β were mock-infected or infected with ALSV. At 24 or 48 hpi, the mRNA levels of host *IFIT2*, *IFIT3*, and *IRAV* genes in cells were examined using qPCR, with *GAPDH* serving as the internal reference control. **(G)** HEK293T cells were co-transfected with an ISRE-luc plasmid and a control plasmid pGL4.74. At 24 h post-transfection (hpt), cells were infected with ALSV and treated with IFN-β for 24 h. Cells were harvested, and luciferase activity was measured. **(H and I)** HEK293T cells treated with IFN-β were mock-infected or infected with ALSV. At 48 hpi, cells were analyzed by immunoblotting with the indicated antibodies. Gray-scale statistical analysis of pSTAT2 relative to STAT2, STAT2 relative to GAPDH (H), and IFIT1, IFIT3, IRAV relative to GAPDH (I) are displayed on the right. Statistical analysis was performed using one or two-way ANOVA with multiple comparison correction (\**P* < 0.05, \*\**P* < 0.01, \*\*\**P* < 0.001, and ****P < 0.0001).

Efficient IFN-I antagonism is of particular importance for all vector-borne flaviviruses, and various strategies for evading IFN response have been demonstrated in viruses such as DENV and ZIKV (Grant et al., 2016). To explore whether ALSV antagonizes the IFN-I response, we treated HepG2 cells with IFN-β and subsequently mock-infected or infected with ALSV. The qPCR results revealed that ALSV infection inhibited the expression of ISGs, including *IFIT2*, *IFIT3* and *IRAV*, especially at 48 hpi (Fig. 2 F). In HEK293T cells, we also found that ALSV infection suppressed IFN-β induced ISRE promoter activity (Fig. 2 G) and reduced the protein levels of phosphorylated STAT2 (Tyr689), and total STAT2, IFIT1, IFIT3, IRAV induced by IFN-β (Fig. 2, H and I). These findings collectively demonstrated that ALSV, like other flaviviruses, has the capability to antagonize the IFN-I response.

### ALSV NSP1 antagonizes IFN-I signaling by degrading STAT2 through an autophagy pathway

To investigate which viral proteins of ALSV regulate IFN-β-induced downstream signaling, we subcloned ALSV genes into a Flag-tagged vector, encompassing nonstructural genes NSP1-2, and structural genes VP1a, VP1b, and VP2-4 (Fig. 3 A) (Zhao et al., 2022). We examined the individual effects of ALSV proteins on activation of ISRE promoter, and found that NSP1 and VP4 significantly inhibited IFN-β-mediated ISRE promoter activation (Fig. 3 B). To further confirm NSP1 can inhibit IFN-I signaling, we transfected A549 cells with NSP1 or vector plasmid. At 24 hours post-transfection (hpt), cells were treated with or without IFN-β for 12 h. QPCR results demonstrated that NSP1 inhibited the mRNA levels of *IFIT1*, *IFIT2*, *IFIT3* and *IRAV* (Fig. 3 C). In HEK293T cells, we also observed that NSP1 suppressed IFN-β induced ISGs expression in a dose-dependent manner (Fig. S1 A), and reduced the protein levels of IFIT1 and IFIT3 induced by IFN-β (Fig. 3, D and E). These findings provide strong evidence that ALSV NSP1 inhibits the expression of ISGs induced by IFN-β.

**Figure 3.**
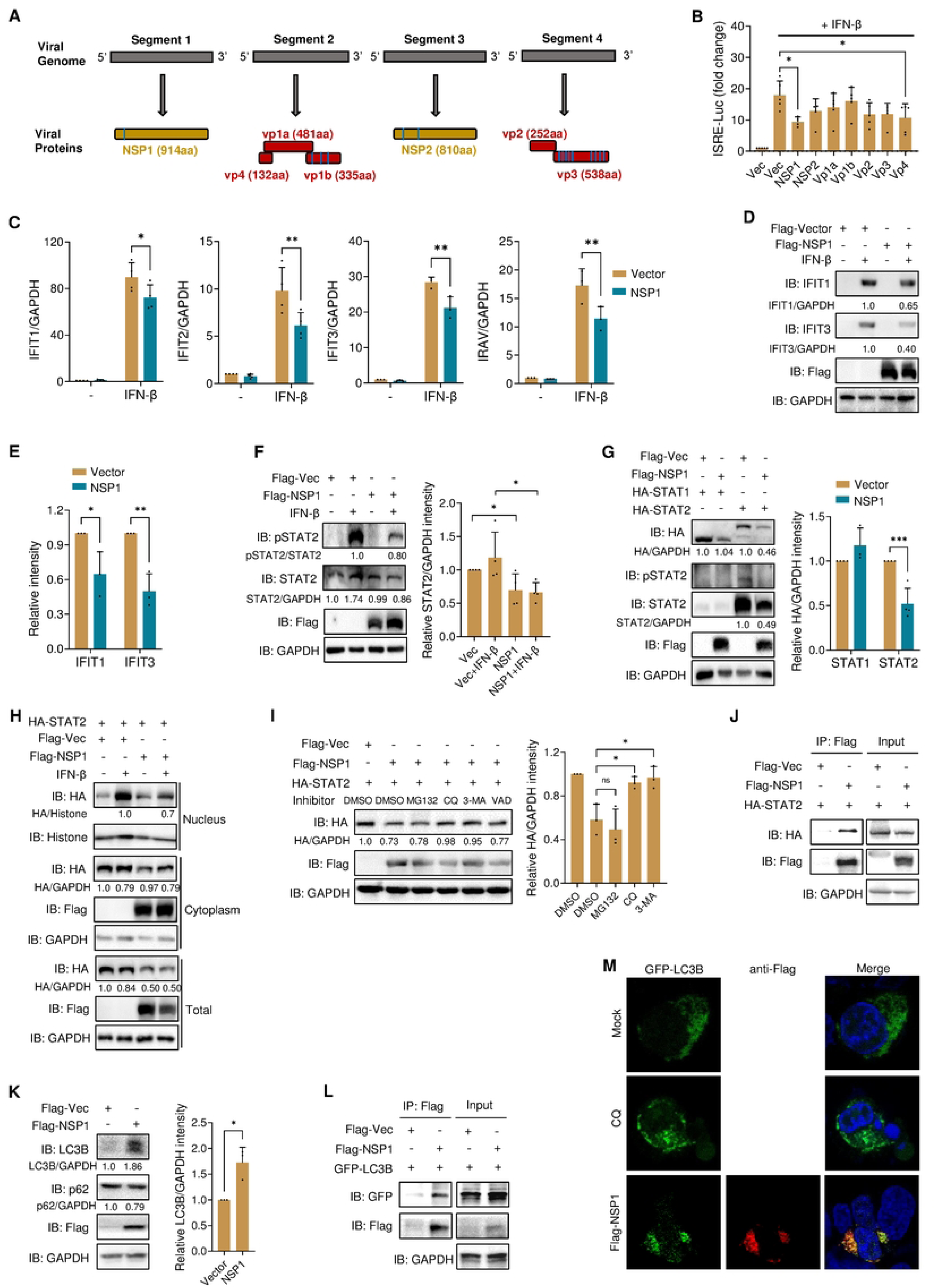
ALSV NSP1 protein antagonizes IFN-I signaling by mediating STAT2 degradation in an autophagy pathway. **(A)** Schematic diagrams of the ALSV genome, comprising four segments that encode structural and nonstructural proteins. The transmembrane region is indicated by the blue box. **(B)** HEK293T cells were transfected with an ISRE reporter plasmid, a control plasmid, and plasmids expressing ALSV proteins. At 24 hours post-transfection (hpt), cells were treated with IFN-β for 12 h, and luciferase activity was measured. **(C)** A549 cells cultured in 24-well plates were transfected with NSP1 or vector plasmid (0.25 μg/well). At 24 hpt, cells were treated with or without IFN-β for 12 h, and the mRNA expression of *IFIT1*, *IFIT2*, *IFIT3*, and *IRAV* was examined using qPCR. **(D)** HEK293T cells were treated as in (C). At 24 hpt, cell lysates were analyzed by immunoblotting with anti-IFIT1, IFIT3, and Flag antibodies, with GAPDH as a loading control. **(E)** The gray-scale statistical analysis of IFIT1 and IFIT3 relative to GAPDH of lines 2 and 4 in (D) is analyzed. **(F)** HEK293T cells were transfected with Flag-tagged NSP1 or vector plasmid. At 24 hpt, cells were treated with or without IFN-β for 12 h, and cell lysates were analyzed by immunoblotting. Gray-scale statistical analysis of STAT2 relative to GAPDH is displayed on the right. **(G)** HEK293T cells were co-transfected with NSP1 and STAT1 or STAT2 expressing plasmids. At 24 hpt, cell lysates were analyzed by immunoblotting. Gray-scale statistical analysis of STAT1 and STAT2 relative to GAPDH is displayed on the right. **(H)** HEK293T cells were co-transfected with NSP1 and STAT2 plasmids. At 24 hpt, cells were treated with or without IFN-β for 30 min, then subjected to cell fractionation assay by immunoblotting. **(I)** HEK293T cells were co-transfected with NSP1 and STAT2 plasmids. At 24 hpt, cells were treated with the inhibitors MG132, CQ, 3-MA, and VAD. After 6 h, cell lysates were analyzed by immunoblotting. Gray-scale statistical analysis of STAT2 relative to GAPDH is displayed on the right. **(J)** HEK293T cells were co-transfected with HA-STAT2 together with NSP1 or vector plasmids. At 24 hpt, cells were treated with CQ for 6 h, and subjected to anti-Flag immunoprecipitates and analyzed by immunoblotting. **(K)** HEK293T cells were transfected with NSP1 or vector plasmid. At 24 hpt, cell lysates were analyzed by immunoblotting with LC3B, p62, and Flag antibodies. Gray-scale statistical analysis of LC3B relative to GAPDH is displayed on the right. **(L)** HEK293T cells were transfected with Flag-NSP1 and GFP-LC3B plasmids. At 48 hpt, anti-Flag immunoprecipitates were analyzed by immunoblotting. **(M)** GFP-LC3 dot formation in HepG2 cells transiently transfected with GFP-LC3B and either left untransfected (Mock) or transfected with Flag-NSP1 for 48 h or treated with CQ for 16 h. Nuclei were counterstained with DAPI. Statistical analysis was conducted using one or two-way ANOVA with multiple comparison correction (\**P* < 0.05, \*\**P* < 0.01, and \*\*\**P* < 0.001).

We then delved into the mechanism through which NSP1 inhibits IFN-I signaling. To investigate whether NSP1-mediated IFN-I antagonism functions upstream or downstream of transcription factors STAT1/STAT2, we assessed the protein levels and activity of STAT1/2 in HEK293T cells expressing NSP1. Immunoblotting revealed that NSP1 reduced IFN-β induced total STAT2 and its phosphorylation, regardless of the presence or absence of IFN-β (Fig. 3, F and G). However, it did not affect total or phosphorylated STAT1 (Fig. S1 B). In addition, the nuclear and cytoplasmic extraction and the IFA analysis showed that the expression of NSP1 prevented IFN-β mediated nuclear translocation of STAT2 (Fig. 3 H and Fig. S1 C). Taken together, our findings reveal that ALSV NSP1 antagonizes IFN-I signaling by targeting STAT2.

The reduction of STAT2 induced by NSP1 may result from the inhibition of protein synthesis or the promotion of protein degradation. Both luciferase and qPCR results revealed NSP1 did not inhibit STAT2 promoter activity and mRNA expression, which were consistent with those observed for STAT1 (Fig. S1, D and E), demonstrating that NSP1 did not affect STAT2 synthesis. Then, we explored whether NSP1 induces STAT2 degradation through specific pathways. HEK293T cells were co-transfected with NSP1 and STAT2 plasmids, followed by treatment with DMSO, the proteasome inhibitor MG132, the autophagy inhibitors CQ and 3-MA, or the pan-caspase inhibitor Z-VAD-FMK, which revealed that CQ and 3-MA reversed NSP1-induced STAT2 reduction (Fig. 3 I), indicating that NSP1 degraded STAT2 through an autophagy pathway. Furthermore, the co-immunoprecipitation (Co-IP) results showed that NSP1 bound to STAT2, especially in the present of CQ, and this binding was independent of IFN-β treatment (Fig. 3 J and Fig. S1 F). We also found that NSP1 induced the expression of autophagosome maker LC3B and increased the LC3-II/LC3-I ratio, while reduced p62 level (Fig. 3 K), and interacted with LC3B (Fig. 3 L). Enhanced autophagic flux was further validated by detecting an increase in GFP-LC3-positive autophagosome formation induced by NSP1, and the co-localization of NSP1 with LC3B and p62 (Fig. 3 M and Fig. S1 G). These results collectively demonstrate that NSP1 degrades STAT2 in an autophagy-dependent pathway.

### NSP1 reduces mitochondrial mass by disrupting mitochondrial dynamics and inhibiting its biogenesis

The membrane (M) protein of SARS-CoV-2 has been shown to reduce mitochondrial mass by inducing mitophagy to suppress IFN-I response, while lipopolysaccharide (LPS) increases mitochondrial mass to promote inflammatory cytokine production, which suggesting mitochondrial mass plays a critical role in immune response (Hui et al., 2021; Yu et al., 2020a). Given that NSP1 reduces the mitochondria mass (Fig. 4 A) (Zhao et al., 2022), we hypothesized that NSP1-induced reduction in mitochondrial mass may be related to IFN-I antagonism. To assess mitochondrial mass, we used MitoTracker-Red (MTR), a fluorescent probe independent of mitochondrial Δψm. We conducted IFA and flow cytometry analysis to measure the MTR fluorescence intensity in various cells expressing Flag- or GFP-tagged NSP1, which consistently demonstrated that NSP1 reduced mitochondrial mass (Fig. 4, B and C; and Fig. S2, A and B). Furthermore, NSP1 dose-dependently reduced the levels of mitochondrial marker proteins, such as COXIV, TOM20, and TIM23, while not affecting the endoplasmic reticulum marker calnexin (Fig. 4 D). Additionally, NSP1 led to a time-dependent decrease of STAT2, TOM20, and TIM23, accompanied by an increase in the LC3-II/LC3-I ratio and a decrease in p62 (Fig. 4 E). These findings collectively confirm that NSP1 induces mitochondrial mass reduction and mitophagy.

**Figure 4.**
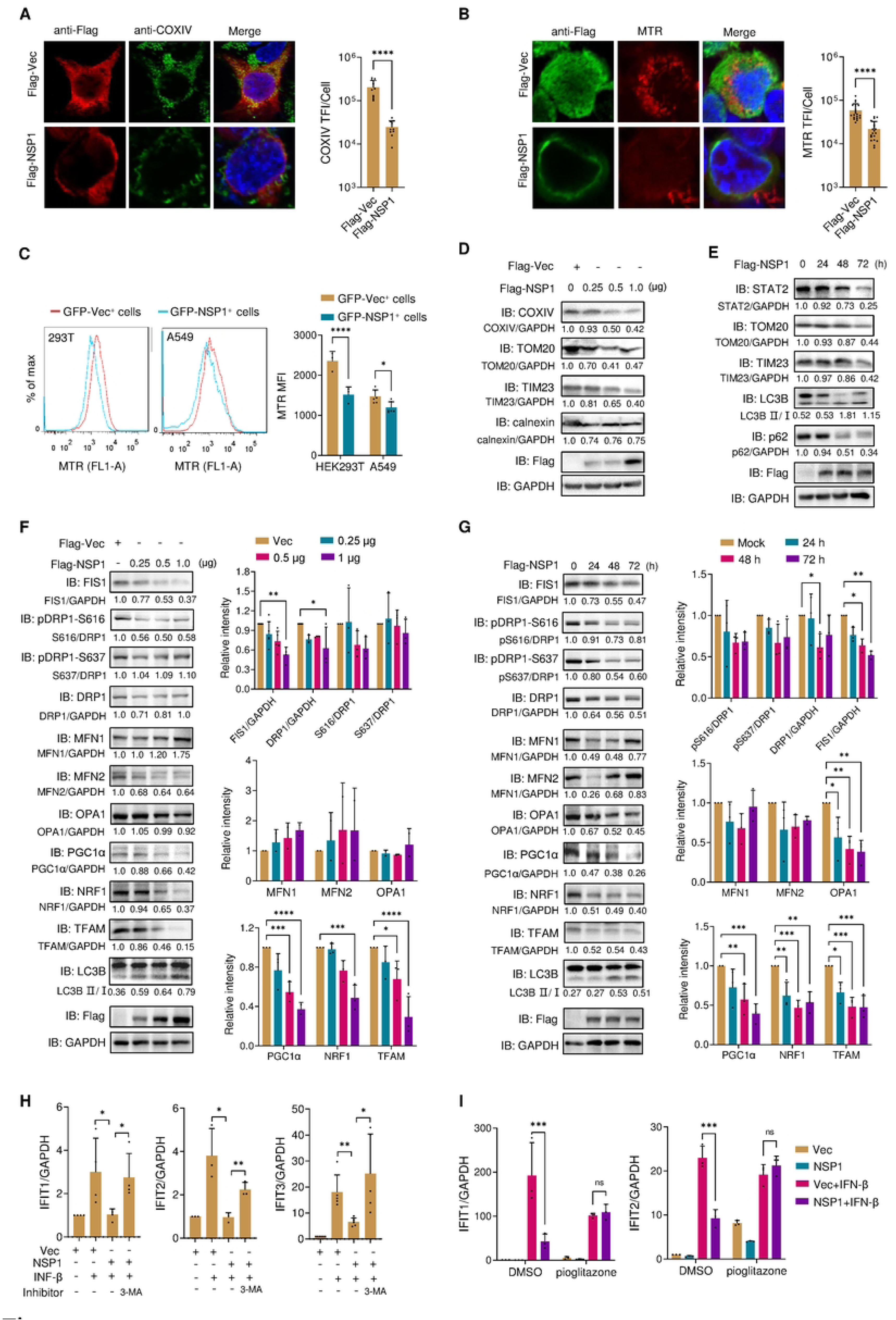
NSP1 reduces mitochondrial mass by disrupting mitochondrial dynamics and inhibiting its biogenesis. **(A and B)** HEK293T cells were transfected with Flag-tagged NSP1 or vector plasmid. At 24 hours post-transfection (hpt), cells were subjected to immunofluorescence with anti-Flag and COXIV antibodies (A) or stained with MitoTracker-Red (MTR) (B). The total fluorescence intensity (TFI) of COXIV or MTR per cell was analyzed (n ≥ 10 cells). **(C)** HEK293T or A549 cells were transfected with NSP1 with an unfused GFP tag. At 48 hpt, cells were stained with MTR and detected by flow cytometry. The mean fluorescence intensity (MFI) of MTR was analyzed in GFP^+^ cells (n ≥ 3). **(D)** HEK293T cells were transfected with escalating doses of NSP1 plasmid. At 24 hpt, cell lysates were analyzed by immunoblotting with the indicated antibodies. **(E)** HEK293T cells were transfected with NSP1 and collected at the indicated times. Cell lysates were analyzed by immunoblotting. **(F)** HEK293T cells were transfected with the escalating doses of NSP1. At 24 hpt, cell lysates were analyzed by immunoblotting. Gray-scale statistical analysis is displayed on the right. **(G)** HEK293T cells were transfected with NSP1 and collected at the indicated times. Cell lysates were analyzed by immunoblotting. Gray-scale statistical analysis is displayed on the right. **(H)** HEK293T cells were transfected with NSP1 or vector. At 24 hpt, cells were treated with or without IFN-β and 3-MA for 12 h, and qRT-PCR measured the mRNA levels of *IFIT1*, *IFIT2*, and *IFIT3*. (*I*) HEK293T cells were transfected with NSP1 or vector. At 24 hpt, cells were treated with or without IFN-β and pioglitazone (1μM) for 16 h, and qRT-PCR measured the mRNA levels of *IFIT1* and *IFIT2*. Statistical analysis was conducted using one or two-way ANOVA with multiple comparison correction (\**P* < 0.05, \*\**P* < 0.01, \*\*\**P* < 0.001, and \*\*\*\**P* < 0.0001).

The regulation of mitochondrial mass, which encompasses mitochondrial dynamics, mitophagy, and mitochondrial biogenesis, constitutes a pivotal mechanism for coordinating the structure and function of mitochondria (Chang et al., 2022). Mitochondrial dynamics is the essential physiological process of mitochondrial fusion and fission. Of which, mitochondrial fusion can facilitate the metabolites rapid exchange and complement impaired mitochondria to enhance their functionality, which is controlled by the outer membrane fusion protein, mitofusins 1 and 2 (MFN1 and 2), and the inner membrane fusion protein, optic atrophy 1 (OPA1) (Archer, 2013; Kabra and Jastroch, 2023). On the other hand, mitochondrial fission involves the division of mitochondria into two new organelles and is mediated by conserved dynamin family GTPases: mitochondrial fission 1 protein (FIS1) and dynamin-related protein 1 (DRP1) (Kabra and Jastroch, 2023). Subsequently, damaged mitochondria are targeted for degradation through mitophagy, a specific autophagic process responsible for eliminating dysfunctional mitochondria (Zhang et al., 2022). To elucidate the underlying mechanisms through which NSP1 reduces mitochondrial mass, we investigated the expression of proteins involved in mitochondrial dynamics, and found that NSP1 reduced the levels of fission-related proteins FIS1 and DRP1 in a dose-dependent manner, without affecting DRP1 phosphorylation or the fusion-related proteins (Fig. 4 F). Furthermore, NSP1 decreases the levels of FIS1, DRP1, and OPA1 in a time-dependent manner, while other mitochondrial dynamics-related proteins remain unaffected (Fig. 4 G). These results suggest that NSP1 disrupts mitochondrial dynamics, leading to subsequent mitophagy.

A reduced mitochondrial population can trigger mitochondrial biogenesis, a process is controlled by the transcriptional factors peroxisome proliferator activator receptor gamma-coactivator 1α (PGC-1α), nuclear respiratory factor 1 (NRF1), and mitochondrial transcription factor A (TFAM). Previous studies have shown that STAT2 promotes LPS-induced mitochondrial mass increase through the transcriptional activation of mitochondrial biogenesis (Yu et al., 2020a). Given that NSP1 can degrade STAT2 (Fig. 3), we speculate that NSP1 might inhibit mitochondrial biogenesis. As expected, our experiments revealed that NSP1 indeed inhibited the expression of PGC-1α, NRF1 and TFAM (Fig. 4, F and G). Peroxisome proliferator-activated receptors-γ (PPARγ) is a ligand-activated transcription factor that plays a role in regulating PGC1α expression (Jamwal et al., 2021), and the PPARγ agonists such as pioglitazone or rosiglitazone has been shown to stimulate mitochondrial biogenesis and enhance mitochondrial function in conditions like Alzheimer’s disease, Parkinson’s disease, and Huntington’s disease (Corona and Duchen, 2016; de Oliveira Bristot et al., 2019). We further investigate the relationship between NSP1-mediated mitochondrial mass reduction and IFN-I antagonism, and found that the mitophagy-specific inhibitor 3-MA and the PPARγ agonist pioglitazone could reverse NSP1-mediated inhibition of ISGs (Fig. 4, H and I). These results suggest that the imbalance in mitochondrial dynamics and the inhibition of mitochondrial biogenesis contribute to NSP1-induced mitochondrial mass reduction and IFN-I antagonism.

### ALSV reduces mitochondrial mass in a STAT2-dependent manner

Recent studies have highlighted the pivotal role of STAT2 in maintaining mitochondrial homeostasis by promoting mitochondrial fission and biogenesis (Dasgupta et al., 2015; Shahni et al., 2015; Yu et al., 2020a). To explore whether NSP1 reduces mitochondrial mass in a STAT2-dependent manner, we knocked out the STAT2 gene in HEK293T cells (Fig. S3 A). Consistent with previous studies, we observed a reduction in mitochondrial mass in STAT2^-/-^ cells (Fig. S3 B). We transfected GFP-tagged NSP1 or vector plasmids into WT and STAT2^-/-^ cells. After 48 h, cells were stained with MTR and analyzed by flow cytometry. The mean fluorescence intensity (MFI) of GFP-positive cells indicated that NSP1 was no longer able to reduce mitochondrial mass in STAT2^-/-^ cells (Fig. 5 A). Similarly, mitochondrial mass was not decreased in STAT2^-/-^ cells infected with the lentiviral particles expressing GFP-NSP1 (Fig. S3 C). IFA results confirmed that the absence of STAT2 rendered NSP1 unable to further reduce mitochondrial mass (Fig. 5 B). Furthermore, immunoblotting results showed that NSP1 did not reduce the levels of mitochondrial proteins COXIV, TOM20, and TIM23, nor increase the ratio of LC3-II/LC3-I in STAT2^-/-^ cells, suggesting that NSP1 cannot induce stronger mitophagy without STAT2 (Fig. 5 C). Additionally, in STAT2^-/-^ cells, NSP1 did not lead to further reduction of proteins associated with mitochondrial dynamics (FIS1 and DRP1) or mitochondrial biogenesis (PGC1α, NRF1 and TFAM) (Fig. 5 D). These findings collectively suggested that ALSV NSP1 relies on STAT2 to reduce mitochondrial mass.

**Figure 5.**
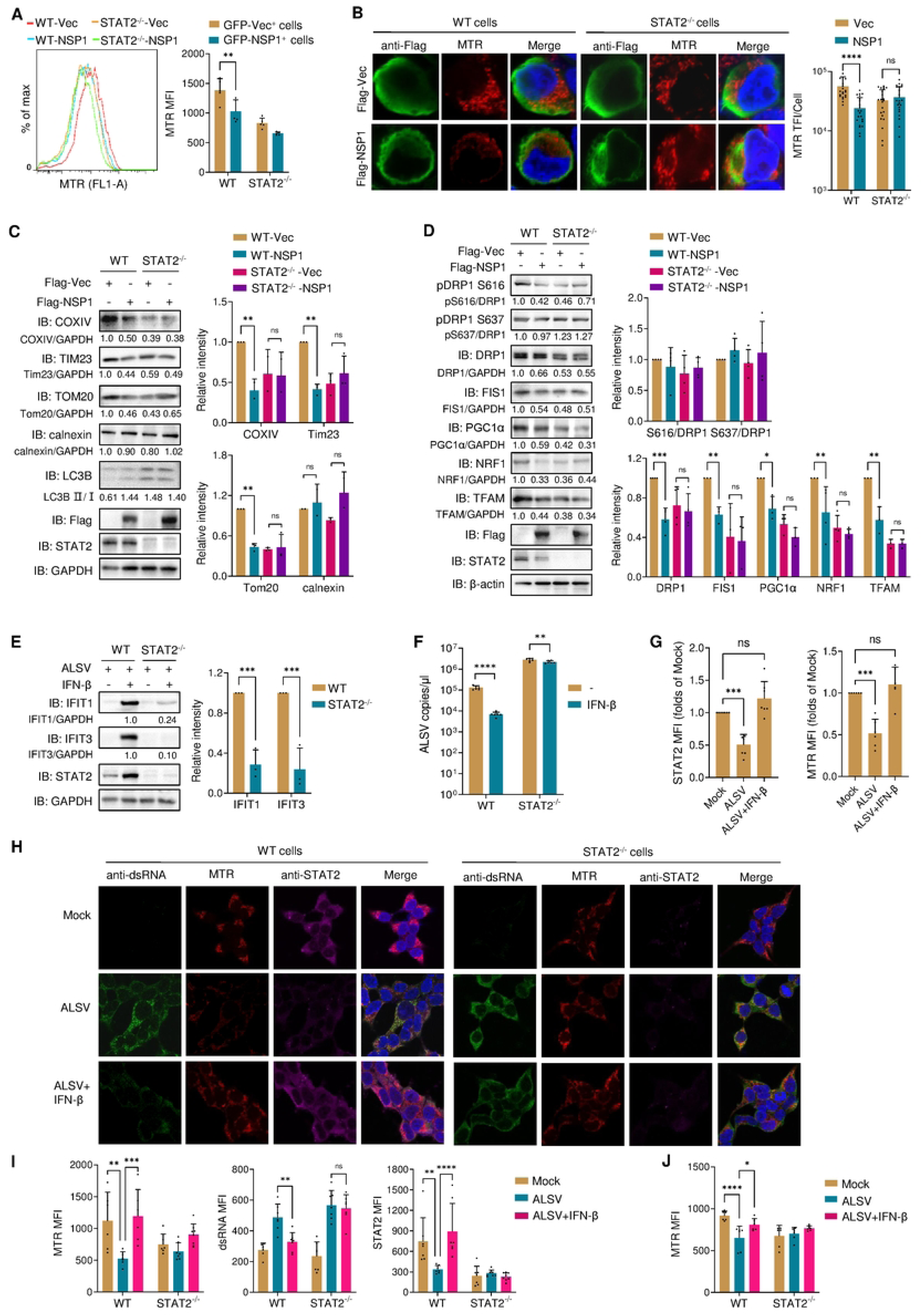
ALSV reduces mitochondrial mass in a STAT2-dependent manner. **(A)** HEK293T wild-type (WT) or STAT2 knockout (STAT2^-/-^) cells were transfected with NSP1 with an unfused GFP tag. At 48 hours post-transfection (hpt), cells were stained with MTR and analyzed by flow cytometry. The MFI of MTR was measured in GFP^+^ cells (n=5). **(B)** WT or STAT2^-/-^ cells were transfected with NSP1 or vector plasmid. At 48 hpt, cells were stained with MTR and anti-Flag antibody. The total fluorescence intensity (TFI) of MTR per cell was measured (n ≥ 20 cells). **(C and D)** WT or STAT2^-/-^ cells were transfected with Flag-NSP1 or vector. At 24 hpt, cell lysates were analyzed by immunoblotting with the indicated antibodies. Gray-scale statistical analysis is displayed on the right. **(E and F)** WT or STAT2^-/-^ cells were infected with ALSV. At 2 hours post-infection (hpi), cells were treated with or without IFN-β for 24 h. Cell lysates were analyzed by immunoblotting with anti-IFIT1, IFIT3, and STAT2 antibodies. Gray-scale statistical analysis of IFIT1 and IFIT3 relative to GAPDH control in lines 2 and 4 is displayed on the right (E). The viral RNA copies in supernatants were determined using TaqMan-qPCR (F). **(G)** A549 cells were infected with ALSV. At 2 hpi, cells were treated with or without IFN-β for 24 h. The cells were stained with anti-dsRNA and STAT2 antibodies. The MFI of STAT2 and MTR per image was measured (n ≥ 5 images). **(H)** HEK293T WT or STAT2^-/-^ cells were infected with ALSV. At 2 hpi, cells were treated with or without IFN-β. Cells were stained with anti-dsRNA and STAT2 antibodies along with MTR. **(I)** The MFI of MTR, dsRNA, and STAT2 per image indicated in (H) was analyzed (n ≥ 6 images). **(J)** WT or STAT2^-/-^ cells were infected with ALSV. At 2 hpi, cells were treated with IFN-β and stained with MTR, then detected by flow cytometry. The MTR MFI was analyzed (n = 4). Statistical analysis was performed using one or two-way ANOVA with multiple comparison correction (\**P* < 0.05, \*\**P* < 0.01, \*\*\**P* < 0.001, and \*\*\*\**P* < 0.0001).

Our observations have revealed ALSV infection suppresses both total STAT2 protein levels and its phosphorylation (Fig. 2 H). In STAT2^-/-^ cells, the ability of IFN-β to induce the expression of ISGs and inhibit ALSV replication is abrogated (Fig. 5, E and F), strongly suggesting that ALSV inhibits IFN-I signaling through its specific targeting of STAT2. Consequently, we proceeded to investigate whether ALSV reduces the mitochondrial mass in a manner dependent on STAT2. Our initial findings indicated that ALSV leads to a decrease in mitochondrial mass, and this reduction is effectively reversed by IFN-β (Fig. 5 G and Fig. S3 D), highlighting the role of IFN-β in mitigating the virus-induced decline in mitochondrial mass. Furthermore, the knockout of STAT2 abolished the inhibitory effect of IFN-β on ALSV replication, and IFN-β no longer counteracted the ALSV-induced loss of mitochondrial mass in the absence of STAT2 (Fig. 5, H and I). These findings were further corroborated by the flow cytometry data, which confirmed that IFN-β failed to eliminate ALSV-mediated reduction in mitochondrial mass in STAT2^-/-^ cells (Fig. 5 J). These data collectively elucidate that ALSV infection also relies on STAT2 to mediate the reduction in mitochondrial mass.

### The interaction map between ALSV NSP1 and STAT2

The structural basis for the suppression of human STAT2 by flavivirus NS5 has been elucidated, revealing a multifaceted interaction between STAT2 and NS5 of ZIKV and DENV. One facet involves the MTase and RdRp domains of NS5 forming a conserved interdomain cleft to prevent the interaction between STAT2 and IRF9 (Wang et al., 2020). In our study, we aimed to examine whether ALSV NSP1 similarly affects the association of STAT2 with IRF9. Co-IP revealed that NSP1 effectively reduces the binding of STAT2 to IRF9 (Fig. 6 A). Another significant interaction involves NS5 binding to key sites F175/R176 located in the coiled-coil domain (CCD) of STAT2 (Wang et al., 2020). This finding raised the intriguing possibility that ALSV NSP1 might interact with STAT2 in a similar manner. As anticipated, our experiments demonstrated that the STAT2 F175A/R176A mutation substantially diminishes the binding of STAT2 to NSP1 (Fig. 6 B). Furthermore, this mutation prevents STAT2 from undergoing degradation by NSP1 (Fig. 6 C), providing compelling evidence that ALSV NSP1 indeed binds to STAT2 through the F175/R176 sites, ultimately leading to STAT2 degradation.

**Figure 6.**
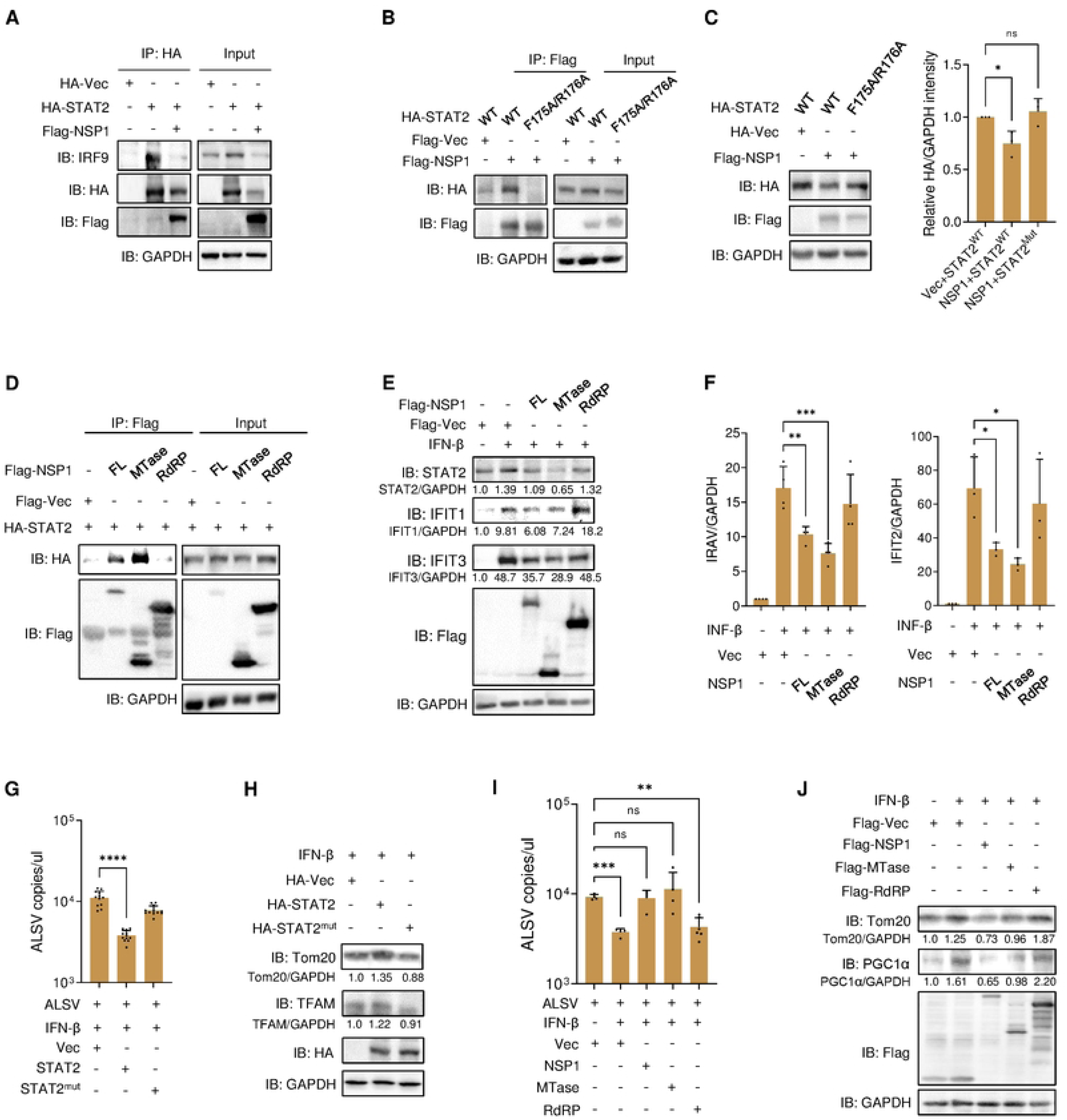
The interaction map between ALSV NSP1 and STAT2. **(A)** HEK293T cells were transfected with HA-STAT2 with or without Flag-NSP1. At 48 hours post-transfection (hpt), anti-HA immunoprecipitates were analyzed by immunoblotting. **(B)** HEK293T cells were transfected with NSP1 or vector along with STAT2 wild-type (WT) or F175A/R176B mutant. At 48 hpt, anti-Flag immunoprecipitates were analyzed by immunoblotting. **(C)** HEK293T cells were transfected with NSP1 or vector along with STAT2 WT or F175A/R176B mutant. At 24 hpt, cell lysates were analyzed by immunoblotting. Gray-scale statistical analysis of HA relative to GAPDH is displayed on the right. **(D)** HEK293T cells were transfected with STAT2 and the indicated truncations of NSP1. At 48 hpt, anti-Flag immunoprecipitates were analyzed by immunoblotting. **(E and F)** HEK293T cells were transfected with plasmids expressing truncations of NSP1 or vector. At 24 hpt, cells were treated with or without IFN-β for 12 h. Cell lysates were analyzed by immunoblotting (E), and the mRNA expression of *IRAV* and *IFIT2* was examined using qPCR (F). **(G and H)** STAT2^-/-^ cells were transfected with STAT2 WT or F175A/R176B mutant. At 24 hpt, cells were infected with ALSV and treated with IFN-β for 24 h. The viral RNA copies in supernatants were determined using TaqMan-qPCR (G), and cell lysates were analyzed by immunoblotting (H). **(I and J)** HEK293T cells were transfected with the indicated truncations of NSP1. At 24 hpt, cells were infected with ALSV and treated with IFN-β for 24 h. The viral RNA copies in supernatants were determined (I), and cell lysates were analyzed by immunoblotting (J). Statistical analysis was conducted using one-way ANOVA with multiple comparison correction (\**P* < 0.05, \*\**P* < 0.01, \*\*\**P* < 0.001, and \*\*\*\**P* < 0.0001).

In the context of ZIKV and DENV NS5, the key sites for interaction with STAT2 have been identified as D734/H855 and D732/L853, respectively (Wang et al., 2020). Given that ALSV NSP1 shares only about 30% amino acid sequence homology with NS5, and notably, the key sites where NS5 binds to STAT2 are not conserved in NSP1 (Fig. S4), we embarked on an investigation to pinpoint which domain of NSP1 is involved in its interaction with STAT2. Remarkably, our findings revealed that the NSP1 MTase domain, but not the RdRp domain of NSP1, is responsible for interacting with STAT2 (Fig. 6 D). Intriguingly, the MTase domain alone is sufficient to induce STAT2 degradation and suppress ISGs expression (Fig. 6, E and F). Furthermore, we transfected STAT2 WT or mutant into STAT2^-/-^ cells, showing that IFN-β was no longer able to inhibit ALSV replication in the presence of STAT2 mutant (Fig. 6 G). Additionally, the expression of the STAT2 mutant in STAT2^-/-^ cells did not lead to an increase in mitochondrial mass (Fig. 6 H). Lastly, our investigation aimed to determined which domain of NSP1 is responsible for promoting ALSV replication by inhibiting IFN-β signaling. Our results conclusively demonstrated that only MTase of NSP1 promoted viral replication and reduces mitochondrial mass (Fig. 6, I and J), providing further substantiation that the MTase domain plays a crucial role in NSP1’s antagonism of the IFN-I response. In summary, our study elucidates that ALSV NSP1 disrupts the association of STAT2 and IRF9, leading to the inhibition of ISGs expression. Moreover, the MTase domain of NSP1 emerges as a key player in this process, shedding light on the intricate mechanisms underlying ALSV evasion of the host immune response.

### ALSV NSP1 exhibits species-specific antagonism of STAT2

Previous studies have demonstrated that the ability of NS5 from DENV and ZIKV to interact with and degrade STAT2 is species-specific, particularly affecting human STAT2 (hSTAT2), but not mouse STAT2 (mSTAT2) (Ashour et al., 2010). This specificity underscores the importance of NS5-mediated IFN antagonism for efficient virus replication. In this context, we were keen to investigate whether ALSV NSP1 could target mSTAT2 for degradation, and found that NSP1 did not reduce the levels of exogenous and endogenous mSTAT2 (Fig. 7, A and B). Furthermore, we explored the interaction between NSP1 and mSTAT2 by performing Co-IPs, showing that NSP1 bound hSTAT2 but not mSTAT2 (Fig. 7 C), and vice versa (Fig. 7 D). Upon re-expressed hSTAT2 or mSTAT2 in STAT2^-/-^ cells, we observed that compared to hSTAT2, NSP1 did not degrade and co-precipitate with mSTAT2 (Fig. 7, E and F). These results conclusively establish that ALSV NSP1 is unable to target mSTAT2 for degradation.

**Figure 7.**
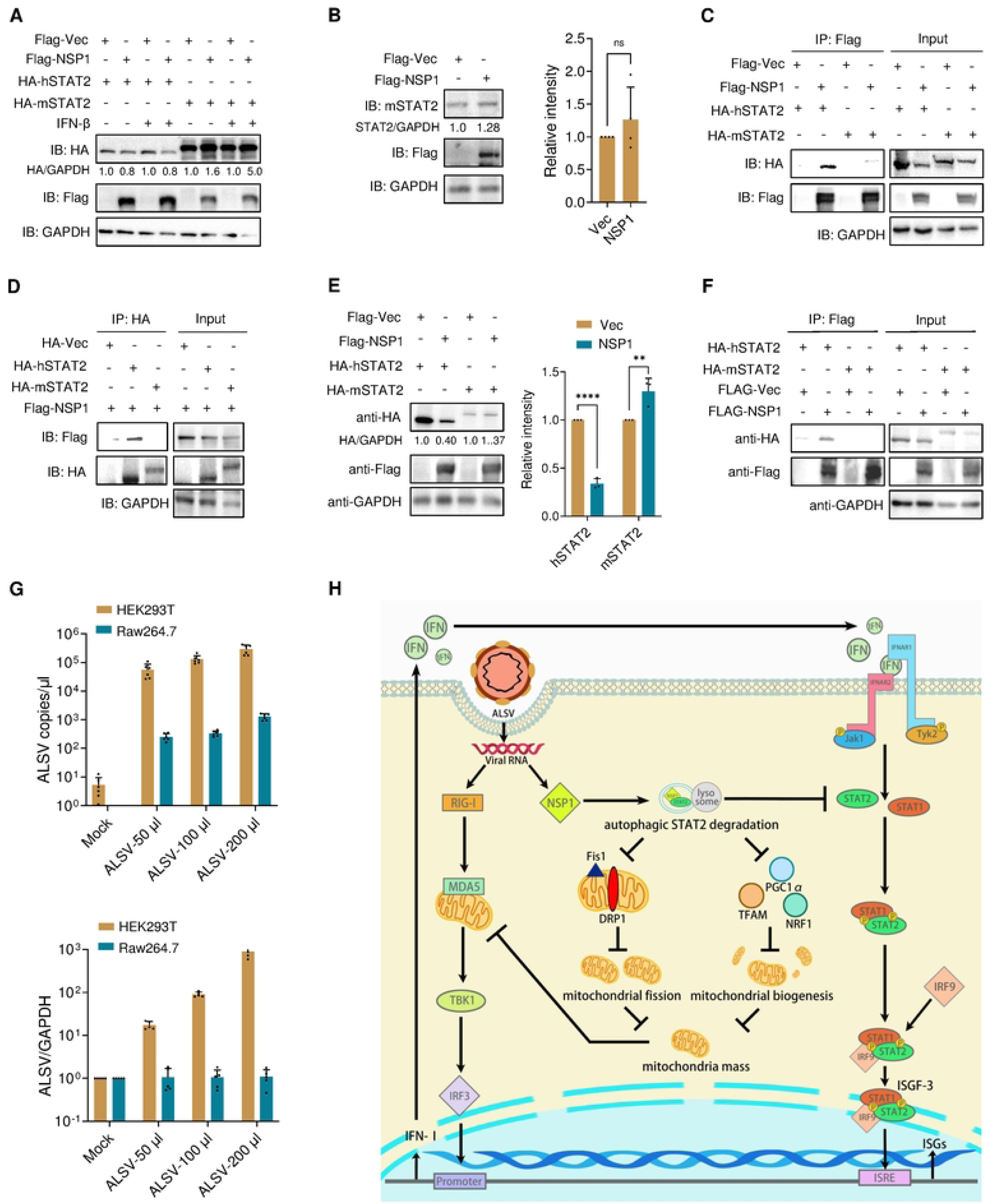
ALSV NSP1 exhibits species-specific antagonism of STAT2. **(A)** HEK293T cells were transfected with NSP1 or vector, along with human STAT2 (hSTAT2) or mouse STAT2 (mSTAT2). At 24 hours post-transfection (hpt), cells were treated with or without IFN-β for 12 h, and cell lysates were analyzed by immunoblotting. **(B)** RAW264.7 cells were co-transfected with NSP1 or vector plasmid. At 24 hpt, cell lysates were analyzed by immunoblotting. Gray-scale statistical analysis of mSTAT2 relative to GAPDH is displayed on the right. **(C and D)** HEK293T cells were transfected with Flag-tagged NSP1 or vector, along with HA-tagged hSTAT2 or mSTAT2. At 48 hpt, anti-Flag (C) or anti-HA (D) immunoprecipitates were analyzed by immunoblotting. **(E and F)** STAT2^-/-^ cells were transfected with hSTAT2 or mSTAT2, along with NSP1 or vector. At 48 hpt, cell lysates were analyzed by immunoblotting, and the gray-scale statistical analysis of HA relative to GAPDH is displayed on the right (E). Moreover, anti-Flag immunoprecipitates were analyzed (F). **(G)** HEK293T and RAW264.7 cells cultured in 24-well plates were infected with the dose-escalated ALSV (50 μl, 100 μl, and 200 μl). At 48 hours post-infection (hpi), the viral RNA copies in supernatants and the viral *S2* mRNA levels in cells were examined using qPCR. **(H)** Schematic representation of the mechanisms underlying ALSV reduces mitochondrial mass via degrading STAT2 to suppress innate immune response. Statistical analysis was conducted using one or two-way ANOVA with multiple comparison correction (\*\**P* < 0.01 and \*\*\*\**P* < 0.0001).

Subsequently, we sought to explore the capability of ALSV to infect mouse-deriving cells. We infected a mouse leukemic monocyte/macrophage cell line RAW264.7 cells with escalating doses of ALSV. At 48 hpi, we assessed the viral RNA copies in supernatants and the viral *S2* mRNA levels in cells using qPCR. Strikingly, our data revealed that ALSV did not replicate in RAW264.7, in contrast to that in HEK293T cells (Fig. 7 G). These findings collectively indicate that ALSV NSP1 exhibits species-specific antagonism of STAT2.

## Discussion

Understanding the pathogenic mechanism of emerging segmented flavivirus serves as a foundational basis for the development of therapeutic drugs and vaccines. In this study, we found that ALSV is sensitive to IFN-β and possesses the capacity to counteract the IFN-I response. Mechanistically, ALSV NSP1 selectively binds to and degrades STAT2 in a species-dependent manner, effectively disrupting the IFN-I signaling pathway. Furthermore, it results in a reduction in mitochondrial mass through perturbing mitochondrial dynamics and inhibiting biogenesis, ultimately leading to the suppression of innate immune response (Fig. 7 H).

The intricate interplay between virus-host interactions that govern induction and evasion of IFN-I response resembles a complex dance, aiming to achieve an optimal balance between virus replication, host disease, and host survival (García-Sastre, 2017). Flavivirus infections are known to trigger the innate immune response by engaging PRRs, subsequently leading to the upregulation of the immune-related gene transcripts and serum IFNs (St John, 2013; Sun et al., 2013). Here, we discovered that ALSV exhibits infection in multiple mammalian cells, and triggers the expression of antiviral genes that play a pivotal role in exerting control over viral replication. On the other hand, viruses must effectively subvert the host’s IFN-I response to replicate or spread to new hosts. In the realm of flaviviruses, various viral non-structural proteins have been identified to interfere with IFN-I response, either by degrading or inhibiting host immune proteins (Aguirre et al., 2017; Dalrymple et al., 2015; Lee et al., 2022; Zhou et al., 2020). Of which, flavivirus NS5 possesses the highest potency and specificity as an IFN antagonist, albeit through virus-specific mechanisms (Grant et al., 2016). WNV NS5 impedes the surface expression of IFNAR1 (Lubick et al., 2015); YFV NS5 relies on IFN treatment to interact with STAT2, thereby preventing STAT2 from binding to ISRE promoter (Laurent-Rolle et al., 2014); DENV NS5 engages the E3 ubiquitin ligase UBR4 to degrade STAT2; ZIKV NS5 targets STAT2 for proteasomal degradation independently of UBR4 (Grant et al., 2016; Morrison et al., 2013). In our study, we unveiled that ALSV NSP1 utilizes autophagy pathway to degrade human STAT2, but not mouse STAT2, which is different from other flaviviruses.

Unlike other STAT family members, STAT2 shows a lower degree of homology between humans and mice (Harrison and Moseley, 2020; Park et al., 1999), which may elucidate the inability of NS5 to degrade murine STAT2. Structural analyses of canonical flavivirus NS5-STAT2 complexes, have unraveled that NS5 MTase and RdRp domains establish a conserved inter-domain cleft that impedes STAT2 association with IRF9. In line with this, our study also found that segmented flavivirus NSP1 binds to STAT2 to inhibit its interaction with IRF9, leading to the inhibition of ISGs. Intriguingly, unlike the NS5 RdRp domain interacts with the key sites F175/R176 of STAT2, NSP1 MTase domain is responsibility for binding to STAT2. This may be caused by the structural differences between NSP1 and NS5 MTase domain, which is unique to segmented flaviviruses (Chen et al., 2023). However, whether NSP1-induced degradation of STAT2 necessitates the presence of the E3 ligase UBR4 remains a subject for further study.

The pivotal link between mitochondria and the innate immune was initially brought to light with the discovery of MAVS, an integral protein localized within mitochondria that plays a crucial role as an adaptor in RIG-I-like receptors signaling (Seth et al., 2005). More recently, other innate immune molecules, such as cGAS, NLRX1, TRAF6, NLRP3, and IRGM, have been functionally associated with mitochondria (Arnoult et al., 2011; Qiu et al., 2023). The accumulating evidences suggest that viruses eliminate critical immune molecules within mitochondria by reducing mitochondrial mass to evade immune response (Mehrzadi et al., 2021; Wang et al., 2021a). For instance, the influenza A virus PB1-F2 protein degrades MAVS by triggering mitophagy (Wang et al., 2021b); coronaviruses, including SARS-CoV-2, promote their replication by altering mitochondrial dynamics and targeting MAVS (Mehrzadi et al., 2021; Shang et al., 2021); ZIKV NS4A induces mitochondrial fission and mitophagy to suppress MAVS-mediated IFN-I response (Lee and Shin, 2023); DENV disrupts mitochondrial biogenesis by downregulating the master regulators PPARγ and PGC1α (Singh et al., 2022). In our present study, we uncovered that ALSV NSP1 disrupts mitochondrial dynamics by reducing the levels of fission-related proteins, FIS1 and DRP1 to induce mitophagy, and inhibits mitochondrial biogenesis through suppressing the expression of PGC-1α, NRF1, and TFAM, ultimately resulting in a decrease in mitochondrial mass. Consequently, we postulate that NSP1 diminishes mitochondrial mass to eliminate immune molecules within mitochondria, such as MAVS and cGAS, thereby inhibiting IFN-I production. Interestingly, promoting mitochondrial mass through inhibiting mitophagy using 3-Methyladenine and enhancing its biogenesis using pioglitazone reversed NSP1-mediated inhibition of ISGs expression.Pioglitazone is a selective PPARγ agonist, and clinically used for the treatment of diabetes (Ha et al., 2023), suggesting that it may become a candidate drug for the clinical treatment of ALSV infection.

Recently, there has been growing recognition of the roles of STATs in mitochondrial function. STAT3 localizes to mitochondria and is implicated in the functioning of OXPHOS complexes to optimize mitochondrial bioenergetics (Gough et al., 2009; Wegrzyn et al., 2009). STAT2 also has been identified within mitochondria (Meier and Larner, 2014). Upon viral infection, STAT2 translocates to mitochondria, where it is targeted for degradation by the viral degradasome formed by viral proteins and MAVS (Goswami et al., 2013). Additionally, Shahni et al. reported that patients harboring clinically asymptomatic STAT2 mutations exhibited severe neurological deterioration following viral infection, which was attributed to STAT2 deficiency, which impeded mitochondrial fission (Dasgupta et al., 2015; Shahni et al., 2015). Furthermore, STAT2 induced by LPS has been found to increase mitochondrial mass by enhancing DRP1 phosphorylation at Ser616. This, in turn, promotes mitochondrial fission and biogenesis, which are essential for the pro-inflammatory differentiation of macrophages (Yu et al., 2020a). Here, we have confirmed that ALSV relied on STAT2 to reduce mitochondrial mass. This discovery enhances our understanding of the intricate relationship between STAT2, mitochondria function, and viral infection. However, the precise mechanism by which NSP1 causes the reduction of mitochondrial mass remains to be explored. It is worth noting that STAT2 could interact with OPA1, but not DRP1 or FIS1 (Fig. S5 A). Moreover, NSP1 does not promote the migration of STAT2 to mitochondria (Fig. S5 B). Further investigations are needed to fully elucidate the specific mechanism involved.

In summary, ALSV is sensitive to IFN-β and possesses the ability to antagonize IFN-I response. At the mechanistic level, ALSV’s NSP1 can selectively bind to and degrade human STAT2 through an autophagy pathway. This degradation directly hampers the expression of ISGs. Furthermore, NSP1-mediated degradation of STAT2 leads to a reduction in mitochondrial mass by perturbing mitochondrial dynamics and suppressing its biogenesis, ultimately resulting in the dampening of antiviral response. In addition, promoting mitochondrial mass through inhibiting mitophagy using 3-Methyladenine and enhancing its biogenesis using pioglitazone can reverse NSP1-mediated antagonism of interferon response, suggesting a potential intervention strategy for ALSV infection. Our findings elucidate the intricate regulatory crosstalk between ALSV and the host’s innate immune response, providing valuable insights into the pathogenesis and intervention strategy for this emerging segmented flavivirus.

## Materials and Methods

### Cells and viruses

HEK293T cells (a human embryonic kidney cell line; ATCC CRL-3216), HepG2 cells (a human hepatoblastoma cell line; ATCC HB-8065), A549 cells (a human lung epithelial cell line; ATCC CCL-185) and Raw264.7 cells (a mouse leukemic monocyte/macrophage cell line; ATCC TIB-71) were grown in Dulbecco’s modified Eagle’s medium (DMEM, high glucose) (Sigma-Aldrich, cat# R8758-500ML), supplemented with 10% fetal bovine serum (FBS) (BBI, cat# E600001) and 1% antibiotics (penicillin and streptomycin, PS) (Sangon, cat# B540732). Vero cells (an African green monkey, Chlorocebus sabaeus, kidney cell line; JCRB0111) were maintained in DMEM (high glucose) supplemented with 10% FBS and 1% PS. STAT2^-/-^ cells, which are STAT2 gene knockout in HEK293 cells, were maintained in DMEM containing 10% FBS, puromycin (2 μg/mL; YEASEN, cat# 60210ES25), and 1% PS. All of the aforementioned cells were cultured at 37 ℃ and 5% CO2 and were routinely tested negative for mycoplasma contamination. To induce an IFN-I response, the target cells were treated with 10 ng/ml recombinant human IFN-β (PeproTech, Cat# 300-02BC) for the specified durations. IRE/CTVM19 cells (a tick, Ixodes ricinus, embryo cell line) were cultured in L15 complete culture medium, which consists of 68% L-15 (Leibovitz) medium, 20% heat-inactivated fetal calf serum, 10% tryptose phosphate broth, 1% L-glutamine and 1% PS, and were maintained at 30 ℃ (Bell-Sakyi et al., 2007).

Alongshan virus (ALSV) was isolated from a patient who had been bitten by a tick in Northeast China (Wang et al., 2019a). Working virus stocks of ALSV were prepared from IRE/CTVM19 cells. In brief, 1 ml of the seed virus was inoculated into IRE/CTVM19 cells. At 1 hpi, an additional 2 ml of fresh L15 complete culture medium was supplemented to the cells. At 7 days post-infection, the culture medium was harvested by centrifuged at 12,000 rpm for 30 min at 4 ℃, and the supernatants were collected to create the working virus stock. All experiments involving infectious ALSV were conducted in strict compliance with Biosafety Level 2 conditions. Manipulations involving both inactivated or non-inactivated ALSV followed the guidelines and regulations set forth by the Chinese authorities regarding dual-use pathogens.

### Plasmid construction

The cDNAs corresponding to the viral protein sequences of ALSV were synthesized by Sangon and subcloned into the XbaI site (YEASEN, Cat# 15033ES76) of 3 × Flag-VR1012 expression vector (BioVector NTCC, Cat# VR1012) using the Hieff Clone Plus One Step Cloning Kit (YEASEN, Cat# 10911) (Zhao et al., 2022). To construct the expression plasmid for ALSV NSP1 MTase (1-280 aa) and RdRp (281-419 aa) domains, the cDNAs were amplified from the Flag-NSP1 template and subsequently cloned into 3 × Flag-VR1012 using Hieff Clone Plus One Step Cloning Kit. For the construction of the plasmid for NSP1 lentiviral transduction, the cDNA of NSP1 were amplified from the Flag-NSP1 template and cloned into the lentiviral vector pWPI (Addgene, Cat# 12254). Plasmid expressing the human STAT2 mutant (HA-STAT2-F175A/R176A) was generated through site-directed mutagenesis PCR using HA-STAT2-Myc (MIAOLING BIOLOGY, Cat# P4105) as a template. To clone mouse STAT2, total RNA was isolated from Raw264.7 cells using the EasyPure® RNA Kit (TransGen, cat# ER101). The synthesized cDNA was PCR amplified using STAT2-specific primers and cloned into the 3 × HA-VR1012 expression vector. Nucleotide sequences were determined by a DNA sequencing service provided by Sangon. Detailed primer sequences are available upon request. The pSTAT1-Luc and pSTAT2-Luc expression plasmids were kindly provided by Dr. Guangyun Tan. All constructed plasmids were transformed into Stbl3 (Thermo Fisher Scientific, Cat# C737303) or Trans5α (TransGen, cat# CD201-01) chemically competent cells, followed by selection on antibiotic-containing LB agar plates.

### ALSV infection

One day before infection, HEK293T cells (5 x 10^4^ cells), STAT2-/- cells (5 x10^4^ cells), A549 cells (5 x 10^4^ cells), HepG2 cells (3 x 10^4^ cells), and Raw264.7 cells (5 x 10^4^ cells) were seeded into 48-well plates. A total of 50 μl of ALSV virus stocks was inoculated and incubated at 37 ℃ in serum-free DMEM (high glucose) for 2 h. The infected cells were subsequently washed twice with PBS and cultured in 300 μl of DMEM (high glucose) containing 2% FBS and 1% PS. For viral RNA quantification, 150 μl of culture supernatant was harvested at the specific timepoints and subjected to Taqman-qPCR using ALSV segment 2 (S2)-specific primers: 5′-GCTT-GTGGTCATCATTATG-3′ (forward), 5′-CTCTGCCACATACTGATG-3′ (reverse), and 5′-CTCTCGTCAGCC-ATACCACCA5-3′ (probe primer). Briefly, viral RNA was extracted from mock- or ALSV-infected cell-culture supernatants using a viral RNA extraction kit (TIANGEN, Cat# DP315), and transcribed into cDNA using the cDNA Synthesis SuperMix (TransGen, cat# AT341). TaqMan-qPCR was performed using the Premix Ex Taq (Probe qPCR) Kit (TaKaRa, cat# RR390L) in an StepOne Plus Real-Time PCR system (Applied Biosystem, USA) with the following cycling conditions: 95 °C for 30 s, followed by 40 cycles of denaturation at 95 ℃ for 5 s and annealing/extension at 60 ℃ for 30 s. The quantitative conversion of virus copy numbers from the Ct value was determined using the formula of (copies)/μl=10^[(Value of Ct-39.01)/-2.895], which was generated by constructing a standard curve with DNA derived from ALSV S2-expressing plasmid stock with a known plasmid concentration. All mock-infected samples exhibited non-singular melting curves, indicating no non-specific amplification, and the values for these samples were set to zero.

Following treatment, the mock- or ALSV-infected cells were collected and used for SYBR-qPCR to quantify the RNA level of viral S2 gene (see “RT-qPCR” section below). Additionally, immunofluorescence analysis was performed on the mock- or ALSV-infected cells to detect viral dsRNA levels.

### Quantitative real-time PCR (RT-qPCR)

Quantitative real-time PCR (qPCR) was performed using the primers listed in the Key Resources Table, following the established protocols (Sui et al., 2021). In brief, after the specified treatment as indicated in Fig. legends, cells were rinsed twice with PBS and then lysed using a lysis buffer. Total RNA was extracted according to the manufacturer’s instructions using the EasyPure RNA Kit. The extracted RNA was subsequently reverse-transcribed into cDNA using the cDNA Synthesis SuperMix (TransGen, cat# AT341). For the qPCR step, Fast SYBR Green Master Mix (Roche, cat# 4913850001) was used, and the reactions were run on a Step-One Plus real-time PCR system. The PCR program started with activation at 95 ℃ for 5 min, followed by 40 cycles of denaturation at 95 ℃ for 10 s and annealing/extension at 60 ℃ for 30 s. The glyceraldehyde 3-phosphate dehydrogenase (GAPDH) gene was employed as the reference gene for normalization. For experiments involving the induction of interferon-stimulated genes (ISGs) measured via qPCR, the target cells were treated with 10 ng/ml IFN-β for the specific durations.

### Dual luciferase reporter assay

For ALSV infection experiment measured via dual luciferase reporter assay (Fig. 2 G), HEK293T cells were seeded in 24-well plates at a density of 2 ×10^5^ per well. They were co-transfected with 250 ng of the IFN-β inducible firefly luciferase reporter plasmid (ISRE-luc) and 50 ng of pGL4.74, which constitutively expresses the renilla luciferase. Transfection was performed using PEI (YEASEN, Cat# 40816ES02) following the manufacturer’s instruction. At 24 hours post transfection (hpt), cells were infected with ALSV and subsequently treated with IFN-β (10 ng/ml) for either 24 or 48 h. Then cells were harvested, and the luciferase activity was measured using the dual-luciferase reporter assay (Promega, cat# E1910). For experiments involving the transfection of viral proteins (Fig. 3 B, 4 F and Fig. S1 A), HEK293T cells cultured in 24-well plates were co-transfected with 500 ng of the indicated plasmids expressing viral proteins, along with 250 ng of ISRE-luc and 50 ng of pGL4.74, all using PEI. At 24 hpt, cells were treated with IFN-β for 12 h and subsequently analyzed for reporter activity. Average firefly luciferase values were normalized to average renilla luciferase values. Mock or empty vector-treated samples without IFN-β treatment was set to 1, and each sample’s luciferase activity was standardized to this value.

### Immunoblotting

For all experiments, HEK293T cells were used, except for Fig. 7 B, where RAW264.7 cells were utilized. HEK293T or STAT2^-/-^ cells were either infected with ALSV or transfected with Flag-tagged NSP1, with or without the indicated plasmids. At 2 hpi or 24 hpt, cells were either mock-treated or treated with 10 ng/ml IFN-β for the specific duration and subsequently analyzed by immunoblotting.

For the STAT2 degradation inhibitor experiment, HEK293T cells transfected with Flag-tagged NSP1 or vector plasmid were treated with the following compounds for 6 h: 10 μM of MG132 (Merck, Cat# 474787), 50 μM of chloroquine (CQ, MCE, Cat# HY-17589A), 10 mM of 3-Methyladenine (3-MA, Selleck, Cat# S2767) or 50 μM of Z-VAD-FMK (Selleck, Cat# S7023). In most cases, cells were lysed for 30 min using a lysis buffer (containing 50 mM Tris-HCl, pH 8.0, 150 mM NaCl, 1% NP-40) supplemented with a protease inhibitor cocktail (Selleck, cat# B14002). To detect phosphorylated proteins, cells were lysed using the same lysis buffer, but with HaltTM protease and phosphatase inhibitor Single-Use cocktail (Thermo Fisher Scientific, cat# 78442). The cell lysates were then centrifuged at 12,000 g for 10 min at 4 ℃, and the protein concentration was determined using Pierce™ BCA Protein Assay Kits (Thermo Fisher Scientific, cat# 23225). The lysates were subsequently reduced with 1 × Protein Loading buffer (TransGen, cat# DL101-02) for 5 min at 95 ℃. For electrophoresis, 20-30 μg of protein for each sample was separated by 8-15% sodium dodecyl sulfate-polyacrylamide gel electrophoresis (SDS-PAGE) and transferred onto PVDF membranes (Millipore, cat# IPVH00010) using Trans-Blot Systems (Bio-rad). Following transfer, the membranes were blocked with 2% BSA (VETEC, cat# V900933) in PBS containing 0.2% Tween-20 (PBST) and then incubated with appropriate primary antibody at 4 ℃ overnight. After washing, the membranes were incubated with HRP-conjugated secondary antibodies, and antibody-antigen complexes were visualized using a chemiluminescence (ECL) substrate (Millipore, cat# WBKlS0100). The grayscale analysis of the bands was performed with ImageJ software.

### Co-immunoprecipitation (Co-IP) assays

For the Co-IP assays, HEK293T cells were transfected with the specified plasmids. At 24 hpt, cells were treated with or without CQ for 12 h prior to the Co-IP procedure. Lysates were collected and centrifuged at 12,000 g for 10 min at 4 ℃ to obtain whole-cell extract. The remaining lysate was then incubated with either anti-Flag M2 Affinity Gel (Sigma-Aldrich, cat# A2220) or anti-HA Affinity Gel (Millipore, cat# E6779) at 4 ℃ overnight to facilitate the immunoprecipitation process. The binding beads were washed several times with lysis buffer and subsequently denatured in 1 x protein loading buffer for 10 min. Finally, the proteins within immunocomplexes and whole-cell extracts were analyzed using immunoblotting with the appropriate antibodies.

### Cells fractionation extraction

The nuclear and cytoplasmic protein extraction process was performed as previously described (Zhao et al., 2021). Briefly, HEK293T cells transfected with Flag-NSP1 and HA-STAT2 plasmids were either treated with or without IFN-β for 30 min. Subsequently, they underwent fractionation extraction using the Nuclear and Cytoplasmic Protein Extraction Kit (Beyotime, cat# P0027) according to the manufacturer’s instructions. The purified nuclear and cytoplasmic proteins, along with total lysates, were then subjected to immunoblotting assays with specific antibodies against Flag, HA, GAPDH, and Histone. For mitochondrial isolation, HEK293T cells were transfected with either Flag-tagged NSP1 or vector plasmid. At 48 hpt, the cells underwent mitochondria extraction using the Cell Mitochondria Isolation Kit (Beyotime, cat# C3601). The isolated mitochondria were subsequently analyzed using immunoblotting.

### Immunofluorescence assays

For viral infection experiments, target cells were seeded onto coverslips in 24-well culture plates amd then infected with ALSV. At 2 hpi, the virus was removed, and the cells were replenished with DMEM medium containing 10 ng/ml IFN-β. After 24 h, cells were fixed with 4% paraformaldehyde (PFA, Biotopped, cat# Top0382) for 30 min. Subsequently, the cells were permeabilized with 0.1% Triton X-100 (YEASEN, cat# 20107ES76) for 15 min and blocking with 1% BSA for 2 h. Following blocking, the cells were stained with anti-dsRNA antibody (SCICONS, Cat# 10010200) and an Alexa Fluor 488-conjugated anti-mouse secondary antibody (Proteintech, Cat# SA00013-1). Finally, nucleic acids within the cells were counterstained with 49, 6-diamidino-2-phenylindole (DAPI) (YEASEN, cat# 40728ES03). To detect STAT2 nuclear transfer, HepG2 cells were transfected with NSP1 or vector plasmid. At 24 hpt, cells were treated with or without IFN-β for 30 min, then they were fixed, permeablized, blocked, and incubated with the anti-Flag and STAT2 antibodies. For the observation of GFP-LC3 dot formation, HepG2 cells expressing GFP-LC3B were either transfected with Flag-NSP1 for 48 h or treated with CQ for 16 h, followed by immunostaining. Cells were visualized using confocal microscopy (FV3000, OLYMPUS), and the acquired images were analyzed with ImageJ software.

### Assessment of mitochondrial mass

MitoTracker® Red CMXRos (MTR, YEASEN, Cat# 40741ES50) is a mitochondrial dye that is independent of mitochondrial membrane potential (Δψm) and is used to monitor mitochondria. One day before infection or transfection, cells, including HEK293T, STAT2^-/-^, A549, and HepG2 cells, were seeded into 24-well plates. The cells were then either infected with ALSV or transfected with the plasmid pWPI-NSP1, which has a GFP tag. After 48 h post-treatment, the cells were incubated with MTR at the final concentration of 100 nM for 30 min at 37°C. Following incubation, the cells were washed three times with PBS for 5 min each and then analyzed by flow cytometry (LSRFortessa, BD Biosciences, MD, USA). The mean fluorescence intensity (MFI) of MTR was analyzed using the FlowJo software. Additionally, the mitochondrial mass is also detected by immunofluorescence assays. Briefly, the cells transfected with the Flag-NSP1 plasmid were fixed and immunostained with anti-Flag and COXIV antibodies after MTR treatment, and the cells infected with ALSV were immunostained with anti-dsRNA and STAT2 antibodies after MTR incubation. Fluorescence images were captured using a confocal microscope. The MFI for infected cells and the total fluorescence intensity (TFI) per cells for transfected cells were quantified using Image J software.

### Lentiviral packaging and transduction

The STAT2 knockout plasmid, pLenti-STAT2-sgRNA (Beyotime, cat# L00231), which simultaneously expresses Cas9, sgRNA for the target gene, and puromycin resistance, were obtained from Beyotime Biotechnology (Shanghai, China). Lentiviral particles were generated by transfecting 8.63 μg of the pLenti-STAT2-sgRNA plasmid, along with 3.1 μg of pMD2.G (Addgene, Cat# 12259) and 5.5 μg of psPAX2 (Addgene, Cat# 12260), into HEK293T cells (6×10^6^ cells) cultured in a 10 cm dish. The transfection was carried out using PEI transfection reagent according to the manufacturer’s protocol. At 48 and 72 hpt, the culture supernatant was harvested, centrifuged, and cell debris was removed by passing it through a 0.45 μm pore size filter (Merck, Cat# SLGVR33RB). Subsequently, lentiviral particles were inoculated into target cells and incubated at 37 ℃ for 48 h. The cells were then selected using culture medium containing 2 μg/mL puromycin (YEASEN, Cat# 60210ES25) for a period of two weeks. After expansion, the expression level of STAT2 was verified through immunoblotting. Simultaneously, the LentiCRISPRv2 plasmid (Addgene, Cat# 52961) was used as an empty vector to establish the control cell line. Polyclonal selected cell populations were used in this study.

To generate lentiviral particles overexpressing ALSV NSP1, HEK293T cells were co-transfected with pWPI-NSP1 or empty pWPI along with pMD2.G and psPAX2 in a ratio of 2.8:1:1.8 using PEI. Viral supernatant was collected at 48 and 72 hpt. The relative infectivity was determined by flow cytometry, measuring the percentage of GFP-positive cells in HEK293T cells infected with serial dilutions of the lentiviral particles for 48 h.

### Quantification and statistical analysis

Data analyses were conducted using Prism 9.0.2 (GraphPad Software). The data are presented as the mean ± standard deviation (SD), and n represents the number of technical replicates. Statistically significant differences were determined by one or two-way analysis of variance (ANOVA) with multiple comparisons correction. Significance is indicated by asterisks, as follows: \**p* < 0.05, \*\**p* < 0.01, \*\*\**p* < 0.001, \*\*\*\**p* < 0.0001, and ns denotes non-significant differences.

### Online supplemental material

Fig. S1 shows ALSV NSP1 protein suppresses IFN-β-induced ISGs expression (related to Fig. 3). Fig. S2 shows ALSV NSP1 reduces mitochondrial mass (related to Fig. 4). Fig. S3 shows ALSV NSP1 reduces mitochondrial mass in a STAT2-dependent manner (related to Fig. 5). Fig. S4 shows sequence alignment of ALSV NSP1 protein with DENV and ZIKV NS5 proteins (related to Fig. 6). Fig. S5 shows the interaction between STAT2 and mitochondria (related to Discussion).

## Acknowledgments

We thank the Tick Cell Biobank (University of Liverpool, UK) that provided the IRE/CTVM19 cell. This work was supported by grant from the National Natural Science Foundation of China (grant numbers 82372250 and 82341105), the Pearl River Talent Plan in Guangdong Province of China (grant number 2019CX01N111), and the Medical Innovation Team Project of Jilin University (grant number 2022JBGS02).

Author contributions

F.J., Y.H., S.F., M.W., L.C., M.P., Ning Liu, L.S., and Y.Z. performed the experiments. W.X., N.L., and L.Z., J.Z., K.Z. prepared experimental materials. Y.Z., F.D., and Z.W. designed the experiments and interpreted the results. Y.Z. wrote the original manuscript. H.G., Nan Liu, Z.H., Z.W., Y.C.Z., and Q.L. revised the manuscript. Q.L. supervised all research. All authors reviewed and proofread the manuscript.

